# Dissociative effects of age on neural differentiation at the category and item level

**DOI:** 10.1101/2023.05.24.542148

**Authors:** Sabina Srokova, Ayse N. Z. Aktas, Joshua D. Koen, Michael D. Rugg

## Abstract

Increasing age is associated with age-related neural dedifferentiation, a reduction in the selectivity of neural representations which has been proposed to contribute to cognitive decline in older age. Recent findings indicate that when operationalized in terms of selectivity for different perceptual categories, age-related neural dedifferentiation, and the apparent age-invariant association of neural selectivity with cognitive performance, are largely restricted to the cortical regions typically recruited during scene processing. It is currently unknown whether this category-level dissociation extends to metrics of neural selectivity defined at the level of individual stimulus items. Here, we examined neural selectivity at the category and item levels using multivoxel pattern similarity analysis (PSA) of fMRI data. Healthy young and older male and female adults viewed images of objects and scenes. Some items were presented singly, while others were either repeated or followed by a ‘similar lure’. Consistent with recent findings, category-level PSA revealed robustly lower differentiation in older than younger adults in scene-selective, but not object-selective, cortical regions. By contrast, at the item level, robust age-related declines in neural differentiation were evident for both stimulus categories. Moreover, we identified an age-invariant association between category-level scene-selectivity in the parahippocampal place area and subsequent memory performance, but no such association was evident for item-level metrics. Lastly, category and item-level neural metrics were uncorrelated. Thus, the present findings suggest that age-related category- and item-level dedifferentiation depend on distinct neural mechanisms.

**Significance Statement:** Cognitive aging is associated with a decline in the selectivity of the neural responses within cortical regions that respond differentially to distinct perceptual categories (age-related neural dedifferentiation). However, prior research indicates that while scene-related selectivity is reduced in older age and is correlated with cognitive performance independently of age, selectivity for object stimuli is typically not moderated by age or memory performance. Here, we demonstrate that neural dedifferentiation is evident for both scene and object exemplars when it is defined in terms of the specificity of neural representations at the level of individual exemplars. These findings suggest that neural selectivity metrics for stimulus categories and for individual stimulus items depend on different neural mechanisms.

## Introduction

Age-related neural dedifferentiation, a reduction in neural selectivity (differentiation) with increasing age, has been proposed to play a role in age-related cognitive decline (Koen & Rugg, 2019; Koen et al., 2020). Evidence suggests that greater neural differentiation at encoding is predictive of better memory performance (Yassa et al., 2011; Berron et al., 2018; Bowman et al., 2019; Koen et al., 2019; Sommer et al., 2019; Srokova et al., 2020), higher scores on psychometric tests of fluid processing (Park et al., 2010; Koen et al., 2019) and might be associated with lower levels of cortical tau deposition in cognitively healthy older adults (Maass et al., 2019). However, despite the apparent functional significance of neural dedifferentiation, the underlying mechanisms remain poorly understood.

Age-related neural dedifferentiation is frequently reported in studies where differentiation is operationalized in terms of the selectivity of neural responses elicited by different categories of visual stimuli (category-level differentiation). Age-related declines in selectivity have almost invariably been reported for scene images (Voss et al., 2008; Carp et al. 2011; Zheng et al., 2018; Koen et al., 2019; Srokova et al., 2020), while effects of age on selectivity for faces and objects are more elusive (for null effects, see Chee et al., 2006; Voss et al., 2008; Zheng et al., 2018; Koen et al., 2019; Payer et al, 2016; Srokova et al., 2020; but see also: Park et al., 2004, 2012; Voss et al., 2008; Zebrowitz et al., 2016). Possible explanations for these mixed findings have been discussed previously (Srokova et al., 2020), but it is remains unknown why neural dedifferentiation is more likely to be observed for scenes than other perceptual categories.

What is the driver of age-related category-level dedifferentiation? One obvious possibility is that it is driven by a decline in the specificity of the neural responses elicited by individual category exemplars. There is, however, no empirical evidence to support this possibility. Whereas category-level dedifferentiation has been extensively documented in prior work, evidence for reduced differentiation at the item level is sparse (see Koen et al., 2020 for discussion). Two prior studies employing multi-voxel pattern analyses reported evidence of an age-related reduction in neural similarity between individual stimuli and their exact repeats (Bowman et al., 2019; Trelle et al., 2019). In another study, which adopted a univariate approach, Goh and colleagues (2010) reported that while younger adults demonstrated attenuated fMRI BOLD responses solely to exact repeats of face images (Barron et al., 2016), older adults demonstrated response reductions both for exact repeats and for items morphed to resemble a previously presented face. This result is suggestive of an age-related broadening of face representations, that is, of reduced neural selectivity. However, contrary to the findings reported in the foregoing studies, two other studies reported null effects of age on item-level neural selectivity (St-Laurent et al., 2019, and, after controlling for category-level effects, Zheng et al., 2018).

In the present study, younger and older adults underwent fMRI as they completed a ‘mnemonic similarity task’ (Stark et al., 2019). During the scanned encoding phase, participants viewed images of objects and scenes. Each novel image was followed either by an exact repetition of the image, a visually similar exemplar, or served as a test item in a subsequent unscanned memory test. Neural differentiation was quantified at both the category and item levels with multivoxel pattern similarity analysis (PSA). On the basis of prior findings (e.g., Koen et al., 2019), we predicted that neural differentiation at the category level would be moderated by age in scene-selective, but not object-selective, cortical regions. Under the assumption that item- and category-level differentiation reflect similar neural mechanisms, we expected to identify analogous age differences in item-level differentiation for scene but not for object images. An open question concerns the patterning of any item-related dedifferentiation effects. Consistent with the findings of Goh et al. (2010), these effects might take the form of an age-related enhancement in neural similarity between novel items and their similar lures, suggestive of a broadening of neural representations. Alternatively, the effects might manifest as attenuated neural similarity between novel items and their repeats, a result consistent with the proposal that neural dedifferentiation at the item level reflects an increase in neural noise (c.f. Li et al., 2001).

## Materials and Methods

### Participants

Twenty-five young and 25 older adults participated in the study. One young and one older adult were excluded from the analyses due to incidental MR findings, resulting in a final sample of 24 young and 24 older adults. Participants were recruited from the University of Texas at Dallas and from the surrounding Dallas metropolitan area and were compensated for their time at a rate of $30/hour and up to $30 for travel. Demographic information and neuropsychological test performance for the final sample are reported in Table 1. Participants were right-handed, had normal or corrected-to-normal vision, and were fluent English speakers before the age of five. None of the participants had a history of neurological or psychiatric disease, substance abuse, diabetes, or current or recent use of prescription medication affecting the central nervous system. All participants undertook a neuropsychological test battery prior to the MRI session, and a set of inclusion and exclusion criteria were employed to minimize the likelihood of including older participants with mild cognitive impairment or early dementia (see below). All participants provided written informed consent before participation in accordance with the requirements of the Institutional Review Board of the University of Texas at Dallas.

**Table 1.** The outcome of the neuropsychological test battery in younger and older adults.

|  | *Younger Adults* | *Older Adults* | *p-value* |
| --- | --- | --- | --- |
| *Age* | 21.54 (3.60) | 68.50 (3.73) |  |
| *Sex (male/female)* | 14/10 | 13/11 |  |
| *Years of Education* | 15.04 (1.63) | 17.21 (2.04) | **< 0.001 \*** |
| *MMSE* | 28.70 (1.22) | 29.12 (0.83) | 0.213 |
| *CVLT Short Delay - Free* | 12.74 (2.49) | 10.58 (3.01) | **0.011 \*** |
| *CVLT Short Delay - Cued* | 12.48 (2.50) | 12.13 (2.46) | 0.628 |
| *CVLT Long Delay - Free* | 12.70 (2.63) | 11.42 (2.89) | 0.120 |
| *CVLT Long Delay - Cued* | 12.82 (2.50) | 12.54 (2.50) | 0.698 |
| *CVLT Recognition Hits* | 15.12 (0.95) | 15.13 (1.15) | 0.765 |
| *CVLT False Alarms* | 1.04 (1.52) | 2.29 (2.69) | 0.057 |
| *F-A-S* | 42.91 (11.79) | 43.71 (11.29) | 0.816 |
| *WMS Logical Memory I* | 28.43 (28.43) | 29.83 (6.60) | 0.466 |
| *WMS Logical Memory II* | 26.09 (6.35) | 27.18 (7.31) | 0.591 |
| *Symbol-digit Substitution Test* | 59.96 (10.81) | 50.75 (8.39) | **0.002 \*** |
| *Trails A (s)* | 22.29 (6.98) | 29.19 (14.34) | **0.042 \*** |
| *Trails B (s)* | 46.51 (15.91) | 55.68 (19.08) | 0.080 |
| *Digit Span* | 18.52 (4.92) | 17.75 (4.44) | 0.575 |
| *Category Fluency Test* | 24.17 (5.33) | 20.54 (3.80) | **0.011 \*** |
| *TOPF* | 49.70 (10.40) | 56.49 (9.16) | **0.023 \*** |
| *Raven's Progressive Matrices I* | 10.91 (1.31) | 9.63 (2.12) | **0.010 \*** |
| *Visual Acuity<sup>1</sup>* | -0.04 (0.09) | 0.13 (0.15) | **< 0.001 \*** |

<sup>1</sup> LogMAR value

### Neuropsychological Testing

Participants completed our laboratory’s standard neuropsychological test battery on a separate day prior to the MRI session. The assessment battery consists of the Mini-Mental State Examination (MMSE), the California Verbal Learning Test II (CVLT; Delis et al., 2000), Wechsler Logical Memory Tests 1 and 2 (Wechsler, 2009), the Symbol Digit Modalities test (SDMT; Smith, 1982), the Trail Making Tests A and B (Reitan and Wolfson, 1985), the F-A-S subtest of the Neurosensory Center Comprehensive Evaluation for Aphasia (Spreen and Benton, 1977), the Forward and Backward digit span subtests of the revised Wechsler Adult Intelligence Scale (WAIS; Wechsler, 1981), The Category Fluency test (Benton, 1968), Raven’s Progressive Matrices List I (Raven et al., 2000), and the Test of Premorbid Functioning (TOPF; Wechsler, 2011). Participants also completed a visual acuity test using ETDRS charts, assessed using the logMAR metric (Ferris et al., 1982; Bailey and Lovie-Kitchin, 2013). Visual acuity was tested with corrective lenses, if prescribed. Participants were excluded prior to the fMRI session if they performed more than 1.5 SD below age norms on two or more non-memory tests, if they performed more than 1.5 SD below the age norm on at least one memory-based test, or if their MMSE score was less than 26. Neuropsychological test scores were missing for one participant (a younger adult male).

### Experimental Materials

Experimental stimuli were presented using PsychoPy v2021.1.3 (Pierce et al., 2019). The study phase was completed inside an MRI scanner and the post-scan memory test was completed post-scan. During the study phase, stimuli were projected onto a translucent screen (41 cm x 25 cm; 1920 x 1080 pixels resolution) placed at the rear of the scanner bore and viewed via a mirror mounted on the head coil (viewing distance approx. 105 cm). The post-scan memory test was administered on a Dell laptop computer equipped with a 17-inch display and a resolution of 1920 x 1080 pixels. All stimuli were presented on a gray background and consisted of images of objects and scenes which were resized to fit inside a frame subtending 256 x 256 pixels.

The critical trials in the study phase comprised 168 scene trials and 168 object trials presented across 9 scanner runs. For a given image category, 48 of the trials (‘first presentation’ trials) were either re-presented (‘exact repeats’, 24 trials) or ‘repeated’ as perceptually similar lures (‘lures’, 24 trials). An additional 72 stimuli belonging to each image category were presented once only, and these items were used as test items in the subsequent memory task (see below). To ensure that neural similarity between the first presentation trial and its corresponding lure or exact repeat was not driven by within-session autocorrelation of the BOLD time series (Mumford et al., 2014), lures and exact repeat trials were always presented in the subsequent scanner run while ensuring that the lag between first and second presentations conformed to a rectangular distribution ranging between 18 and 42 trials (mean = 30). Consequently, the first study run did not contain any repetition or lure trials, and the last run did not contain any first presentation trials; an additional 24 filler trials were randomly interspersed within the first and last runs to ensure that all runs contained an equal number of trials. One hundred and twenty-six null trials, comprising a white fixation cross presented at the center of the display, were randomly interspersed among the critical trials of the study phase. During the test phase, participants viewed a total of 108 images of scenes and 108 images of objects. For a given image category, 36 trials were repetitions of images that the participant had viewed during the study phase (‘target’), 36 trials were images that were perceptually similar to previously viewed study images (‘lure’), and 36 trials were presentations of new images.

The stimulus pool described above was used to create 24 stimulus lists which were assigned to yoked pairs of younger and older adults. For all stimulus lists, the stimuli were pseudorandomized such that participants viewed no more than 3 consecutive trials of the same visual category or trial type, and no more than 2 consecutive null trials.

### Study and Test Phase

A schematic of the study and test tasks, including examples of the experimental stimuli, is illustrated in Figure 1. Participants received instructions for the study phase and completed a short practice run prior to entering the scanner. Each of the 9 scanner runs of the study phase lasted 4 minutes and 38 seconds. A given study trial began with a red fixation cross presented in the middle of the screen for 500 ms, followed by an image of an object or a scene for 2 seconds, and then a white fixation cross for an additional 2 seconds. Participants had a total of 4 seconds following the onset of the image to make ‘indoor / outdoor’ judgements on the presented scene or object. The indoor and outdoor judgements were mapped to the right and middle finger of the right hand with finger assignment counterbalanced across participants. Responses were made using a scanner compatible button box.

**Figure 1.**
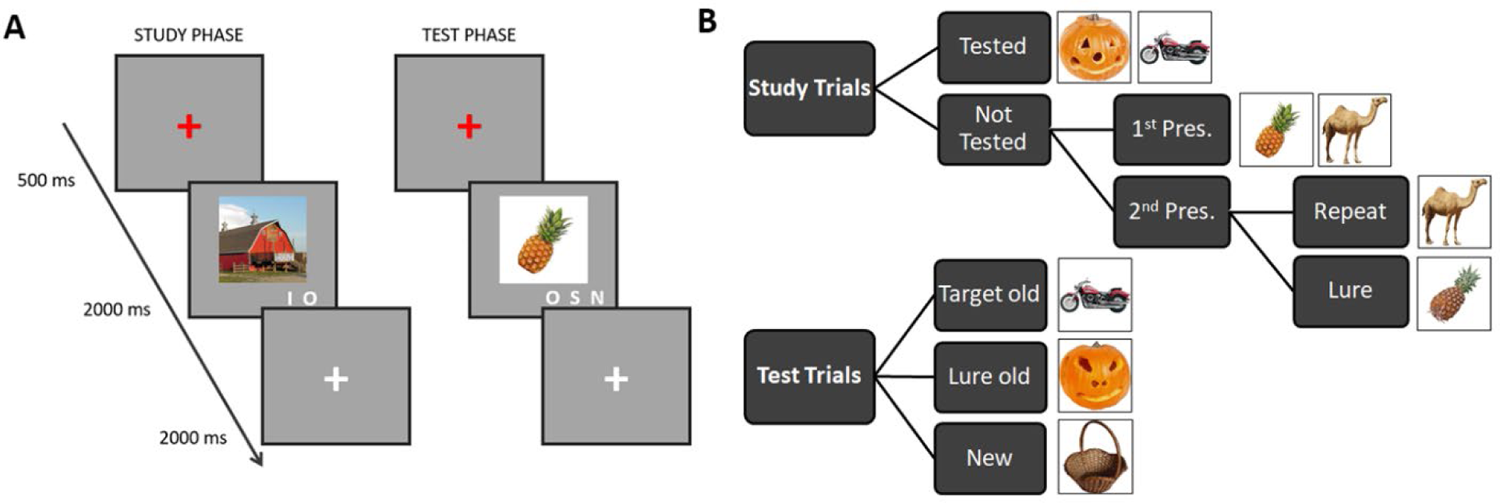
**(A)** An illustration of representative trials from the study and test phases and **(B)** a schematic illustrating the trial types. Study trials were categorized into 5 trial types: tested trials (later employed in the retrieval task), first presentation trials (to be followed by an exact repeat or by a perceptually similar lure in the subsequent scanner run), and second presentation trials (their exact repeats and similar lures). At test, trials were binned into three trial types: target items (previously studied exemplars), lure items (similar to a studied exemplar), or new items. At study, participants made ‘indoor/outdoor’ judgements, and at test they made ‘old/similar/new’ judgements.

The instructions and a practice run for the test phase were administered immediately following the MRI session. The test phase consisted of 2 blocks lasting approximately 8 minutes each. Each trial of the test phase began with a red fixation cross for 500 ms, followed by the test image for 2 seconds and a white fixation cross for an additional 2 seconds, thus providing a 4 second response window. Participants were instructed to indicate whether the test image was either the same as one they had viewed at study (‘old’), similar to an image they had viewed at study (‘similar’), or a completely new exemplar (‘new’). The three response alternatives were mapped onto the index, middle, and ring fingers of the right hand with finger assignment counterbalanced across participants.

### Online Similarity Rating Task

As discussed in the introduction, one of the aims of the present study was to determine whether metrics of item-level neural differentiation for scene and object exemplars are differentially impacted by age. To ensure that any potential category effects did not arise merely because of categorical differences in visual similarity between items and their similar lures, we collected similarity ratings for pairs of perceptually similar images from 210 online raters. The similarity rating task was programmed in JavaScript using PsychoJS and hosted on Pavlovia.org. Participants were recruited through Prolific.co, compensated $15/h, and provided informed consent in accordance with the requirements of the Institutional Review Board at the University of Texas at Dallas.

We collected ratings on a total of 86 scene and 86 object perceptually similar triplets. The triplets were presented as 3 pairs, such that for a given triplet ‘ABC’, the similar images were presented in three trials ‘AB’, ‘BC’, ‘AC’ randomly interspersed within the stimulus list. The stimulus list included an additional 68 perceptually dissimilar image pairs which served as ‘catch trials’ to ensure that participants were paying attention to the stimuli. As a result, the full stimulus list included a total of 516 lure pairs and 68 catch pairs. Given the length of the list, we split it into two lists of 258 lure pairs each while ensuring that all of the similar image pairs belonging to a given triplet were contained within one list. Each participant was administered one list only; thus each image pair was assessed by 105 raters.

Prior to the similarity judgment task, participants completed an instruction phase and a small number of practice trials to familiarize themselves with the procedures and the rating scale. Each critical trial began with a 500 ms duration red fixation cross followed by a 5-second presentation of the image pair and a continuous ‘slider’ rating scale. During these 5 seconds, participants used a computer mouse to place a response marker anywhere along the scale. For the purposes of the analysis of similarity ratings, the rating scale ranged from 0 (not at all similar) to 10 (very similar). The task lasted approximately 30 minutes with a 30 second break provided halfway through.

Figure 2B illustrates across-subject similarity ratings for all object and scene image pairs as well as object and scene catch trials. The similarity ratings for catch trials were robustly lower relative to lure trials, indicating that participants complied with the task instructions. Across all trials, object lure pairs were on average rated as more similar (M = 6.85) than scene lure pairs (M = 6.51, p < 0.001). To ensure that the scene and object lures employed in the fMRI experiment were matched in perceptual similarity, we followed these steps: First, we sorted the lure pairs belonging to each triplet as the ‘most similar pair’, ‘middle pair’, and ‘least similar pair’, according to their across-participant ratings. Next, we selected all pairs from the ‘most similar’ scene group, and all ‘middle pairs’ from the object group, resulting in a total of 86 scene and 86 object pairs. Lastly, we iteratively and randomly selected 60 scene and 60 object pairs until the average across-participant similarity ratings for scenes and objects were equal. The outcome of the matching procedure is illustrated in Figure 2B (see ‘Matched’). The average across-participant ratings for the final 60 lure pairs employed in the fMRI experiment was 6.74 for both objects and scenes.

**Figure 2.**
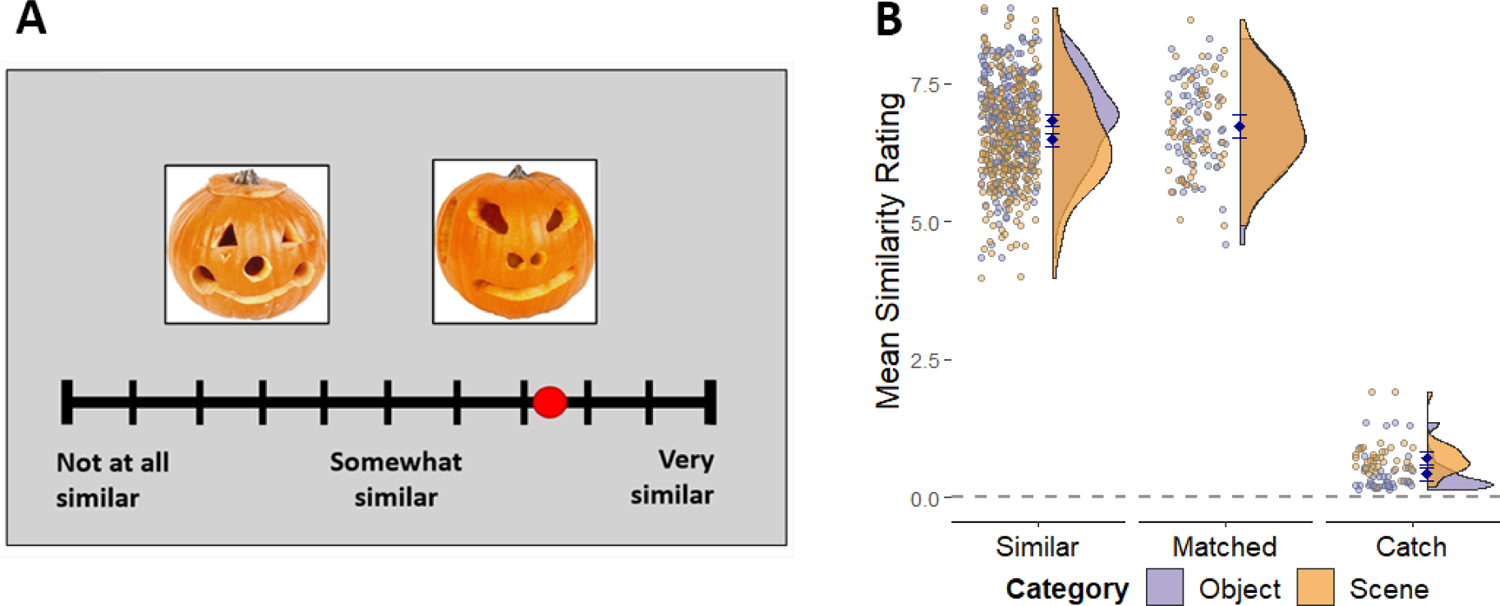
**(A)** An illustration of a single trial in the online similarity rating task. **(B)** Average across-subject similarity ratings for object and scene lures, demonstrating that pairs of scene lures were on average rated as less similar than object lures. Object and scenes did not differ in their similarities following the similarity matching procedure (see main text).

### MRI Data Acquisition and Preprocessing

Functional and structural MRI data were acquired using a Siemens Prisma 3T scanner at the Sammons BrainHealth Imaging Center at the University of Texas at Dallas. The data were acquired with a 32-channel head coil. A whole-brain anatomical scan was acquired with a T1-weighted 3D MPRAGE pulse sequence (FOV = 256 × 256 mm, voxel size = 1 × 1 × 1 mm, 160 slices, sagittal acquisition). Functional data were acquired with a T2*-weighted blood-oxygen-level-dependent echoplanar imaging (EPI) sequence with a multiband factor of 3 (flip angle = 70°, FOV = 220 × 220 mm, voxel size = 2 x 2 x 2 mm, TR = 1.52 ms, TE = 30 ms, 66 slices). A dual echo fieldmap sequence which matched the 3D characteristics of the EPI sequence was acquired at TEs of 4.92 ms and 7.38 ms immediately after the last run of the study phase, resulting in two magnitude images (one per echo), and a pre-subtracted phase image (the difference between the phases acquired at each echo).

The MRI data were preprocessed using Statistical Parametric Mapping (SPM12, Wellcome Department of Cognitive Neurology) and custom MATLAB code (The MathWorks). The functional data were preprocessed in 6 steps. First, we employed the FieldMap toolbox in SPM to calculate voxel displacement maps prior to the fieldmap correction. These maps were calculated using the magnitude and phase difference images acquired in the aforementioned dual-echo fieldmap sequence. Second, SPM’s realign & unwarp procedure was applied, operating in two steps: spatial realignment of the time series registered to the mean EPI image and a dynamic correction of the deformation field using the voxel displacement maps. Third, the functional images were reoriented along the anterior and posterior commissures, then spatially normalized to SPM’s EPI template, and renormalized to an age-unbiased sample-specific EPI template according to procedures standardly employed in our laboratory (see de Chastelaine et al., 2016). Lastly, the functional data were smoothed with a 5 mm FWHM Gaussian kernel.

### Region of Interest Selection

The scene-selective parahippocampal place area (PPA) and the object-selective lateral occipital complex (LOC; Figure 3) were selected as our two *a priori* regions of interest (ROIs), following the analyses reported by Koen et al., (2019). Prior to the ROI selection, the fMRI data were analyzed with a two-stage univariate GLM approach. At the subject-level, all study trials were binned into 10 events of interest (5 events separately for scene and object trials): 1. Trials which went on to be presented in the memory test, 2) first presentation of a to-be-repeated image, 3) first presentation of images which were followed by similar lures, 4) exact repeats, 5) similar lures. Neural activity elicited by the events of interest was modeled with a boxcar function extending over a 2s period coincident with image presentation. The boxcar functions were convolved with a canonical hemodynamic response function (HRF) to estimate the predicted BOLD responses. Additional regressors in the design matrix were trials of no interest (filler trials and trials with missing responses or with responses occurring within 500ms post-stimulus onset), 6 motion regressors reflecting rigid-body translation and rotation, spike covariates regressing out volumes with displacement greater than 1mm or 1° in any direction, and the mean signal of each run. Prior to model estimation, the fMRI time series from each scanner run was concatenated into a single session using the spm_fmri_concatenate function.

**Figure 3.**
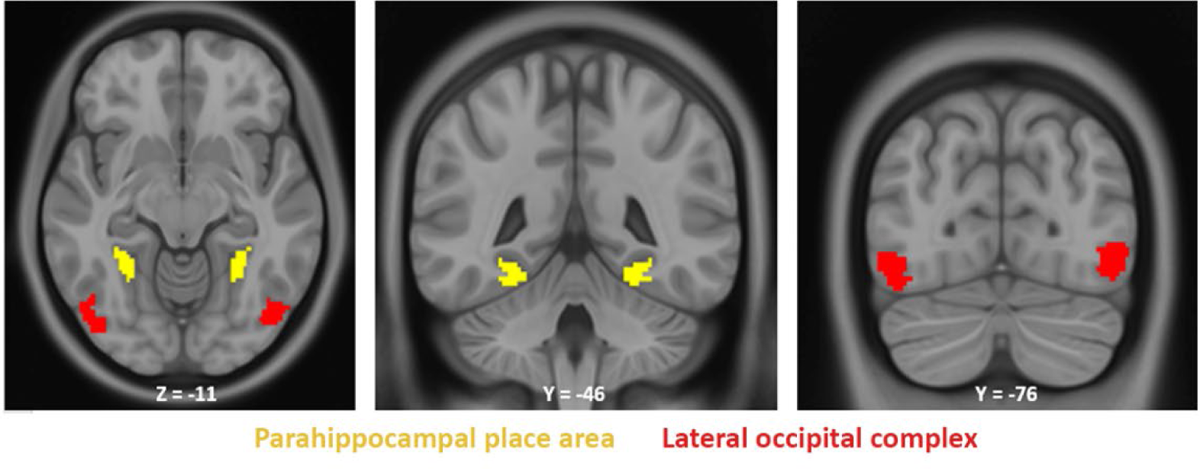
A priori regions of interest (PPA and LOC) for one representative younger/older adult pair illustrated on a T1-weighted ICBM 152 MNI brain.

At the second level, the parameter estimates from the subject-wise GLMs were entered into a group-level mixed factorial ANOVA with factors of age group (2) and events of interest (10). To ensure that the ROIs were derived independently of the to-be-analyzed data and were unbiased with respect to age group, we employed a leave-one-pair-out approach where group-level GLMs were generated by iteratively leaving out a randomly yoked pair of a younger and an older adult (cf. Hill et al., 2021). As a result, ROIs for each held out pair were derived from the data belonging to the remainder of the sample. The ROIs were defined using category-selective contrasts and comprised all voxels that fell within a 10mm radius of the cluster’s peak and were included in anatomical masks provided by the Neuromorphometrics atlas (available in SPM12). The PPA was defined using a scene > object group-level contrast, inclusively masked by the parahippocampal and fusiform gyri (left PPA: M = 147 voxels, SD = 4.5; right PPA: M = 156 voxels, SD = 1.7). The LOC was delineated with an object > scene contrast, restricted by the labels of the inferior and middle occipital gyri (left LOC: M = 286 voxels, SD = 7.5; right LOC: M = 258 voxels, SD = 6.4).

### Multivoxel Pattern Similarity Analysis

For the purposes of the pattern similarity analysis (PSA; Kriegeskorte et al., 2008), the data from the study phase were subjected to a ‘least-squares-all’ GLM (Rissman et al., 2004; Mumford et al., 2014) in which each trial was modeled by a separate 2s boxcar regressor tracking the image presentation. The 6 motion regressors reflecting rigid-body translation and rotation were included as covariates of no interest. Item- and category-level PSA were conducted using approaches similar to those employed in prior work from our laboratory (Koen et al., 2019; Srokova et al., 2020; Hill et al., 2021) and are described in detail below. As noted above, to ensure that category- and item-level similarity metrics were not confounded by within-run autocorrelation effects (Mumford et al., 2014), all correlations were computed between trials belonging to different scanner runs. We also note that the outcomes of all analyses described below were unchanged when visual acuity was employed as a covariate of no interest.

### Category-level PSA

Category-level PSA was operationalized for each ROI (PPA / LOC) and image category (scene / object) by computing the difference between within-category and between-category similarity metrics. Within-category similarity was computed as the average voxel-wise Fisher Z-transformed correlation between a given trial and all trials of the same image category. Between-category similarity was calculated in an analogous fashion – as the average Fisher Z-transformed correlation between a given trial and all trials of the alternate image category. As illustrated in Figure 4, the category-level PSA was performed on novel trials only (i.e., excluding 2^nd^ presentation trials).

**Figure 4:**
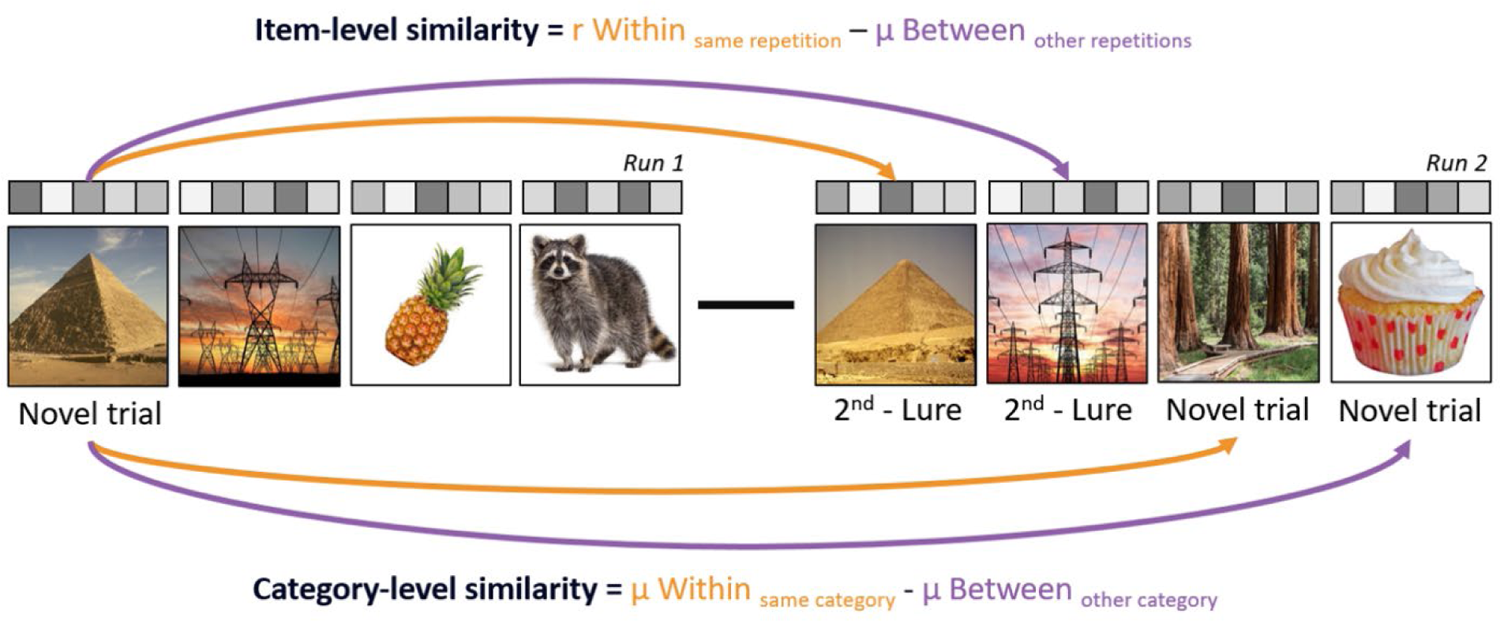
Pattern similarity analysis was employed to quantify neural similarity at the level of individual items and stimulus categories. Item-level similarity was computed as the similarity between a given trial and its repeat or lure minus the average similarity with all other repeats or lures belonging to the same image category. Category-level similarity was computed as the difference between within-category similarity (e.g., average correlation between all scene trials with all other scene trials) and between-category similarity (e.g., average correlation between all scenes with all objects).

### Item-level PSA

Item-level PSA was computed separately for each ROI, image category, and trial type (i.e., exact repeats and similar lures). The item-level similarity metric was computed as the difference between the within-trial and between-trial similarities. Within-trial similarity was computed as the Fisher Z-transformed correlation between a given item and its second presentation (the item’s exact repeat or its similar lure). Between-trial similarity was computed as the average Fisher Z-transformed correlation between a given item and the exact repeat (or lure) trials for every other item belonging to the same category (and belonging to a different scanner run, see Figure 4).

### Whole-brain exploratory PSA

ROIs defined using univariate category-selective contrasts have frequently been employed in prior studies which examined age differences in neural differentiation at the category level (e.g., Koen et al., 2019). However, item- and category-level PSA effects are unlikely to be restricted to regions which exhibit univariate category selectivity. To address this limitation, and following Hill et al., 2021, we supplemented the ROI-based analyses with an exploratory whole-brain PSA conducted across 384 functionally defined cortical parcels comprising the Atlas of Intrinsic Connectivity of Homotopic Areas (AICHA; Joliot et al., 2015). PSA was computed at the item and category levels using the beta-parameters extracted from each AICHA parcel, following the approach described in the preceding paragraphs. We focused these analyses on regions exhibiting positive similarity effects (similarity metrics that were significantly greater than zero) that survived an FDR-adjusted significance threshold of q < 0.05.

### Behavioral Data Analysis

Item recognition performance was assessed separately for object and scene stimuli by computing the difference in the proportion of target trials correctly endorsed ‘old’ (item hits) and the proportion of new trials incorrectly endorsed ‘old’ (false alarms). To quantify discriminability between old targets and lure items, we employed a ‘Target-Lure discriminability’ metric (TLD) – computed as the proportion of old trials correctly endorsed ‘old’ minus the proportion of lure trials which were incorrectly endorsed ‘old’. This metric differs from the more widely employed ‘Lure discrimination index’ (LDI; Stark et al., 2013) which instead indexes discriminability between similar lures and new trials. We consider TLD to be the preferable metric because it is a more direct behavioral correlate of pattern separation – the putative hippocampally mediated process that supports the ability to distinguish between similar inputs (e.g., Yassa et al., 2011).

### Statistical Analyses

Statistical analyses were performed in R studio (R Core Team, 2022). Analyses of variance were performed using the afex package (Singman et al., 2016), with degrees of freedom corrected for non-sphericity with the Greenhouse-Geisser procedure (Greenhouse and Geisser, 1959). T-tests and multiple regressions were performed using the t.test and lm functions in base R. Partial correlations were conducted using pcor. test in the ppcor package (Kim, 2015).

## Results

### Neuropsychological Performance

Neuropsychological test performance is illustrated in Table 1. Young adults outperformed older adults on the following measures: CVLT Free recall (short delay), Raven’s progressive matrices I, category fluency test, SDMT, Trails A, and visual acuity. In contrast, older adults performed better than their younger counterparts on the TOPF.

### Memory performance

Trial proportions binned according to category, trial type, and response type are illustrated in Table 2 and Figure 5A. The estimates of item memory performance (hits – false alarms) were entered into a 2 (age) x 2 (image category) mixed effects ANOVA which revealed a significant effect of category (F_(1, 45)_ = 73.027, p < 0.001, partial-η^2^ = 0.619), reflective of better item memory on object than scene trials. The main effect of category was accompanied by a null effect of age (F_(1, 45)_ = 0.852, p = 0.361, partial-η^2^ = 0.019), and the age-by-category interaction was also not significant (F_(1, 45)_ = 2.482, p = 0.122, partial-η^2^ = 0.052). Thus, there were no age differences between young and older adults in item memory performance.

**Figure 5:**
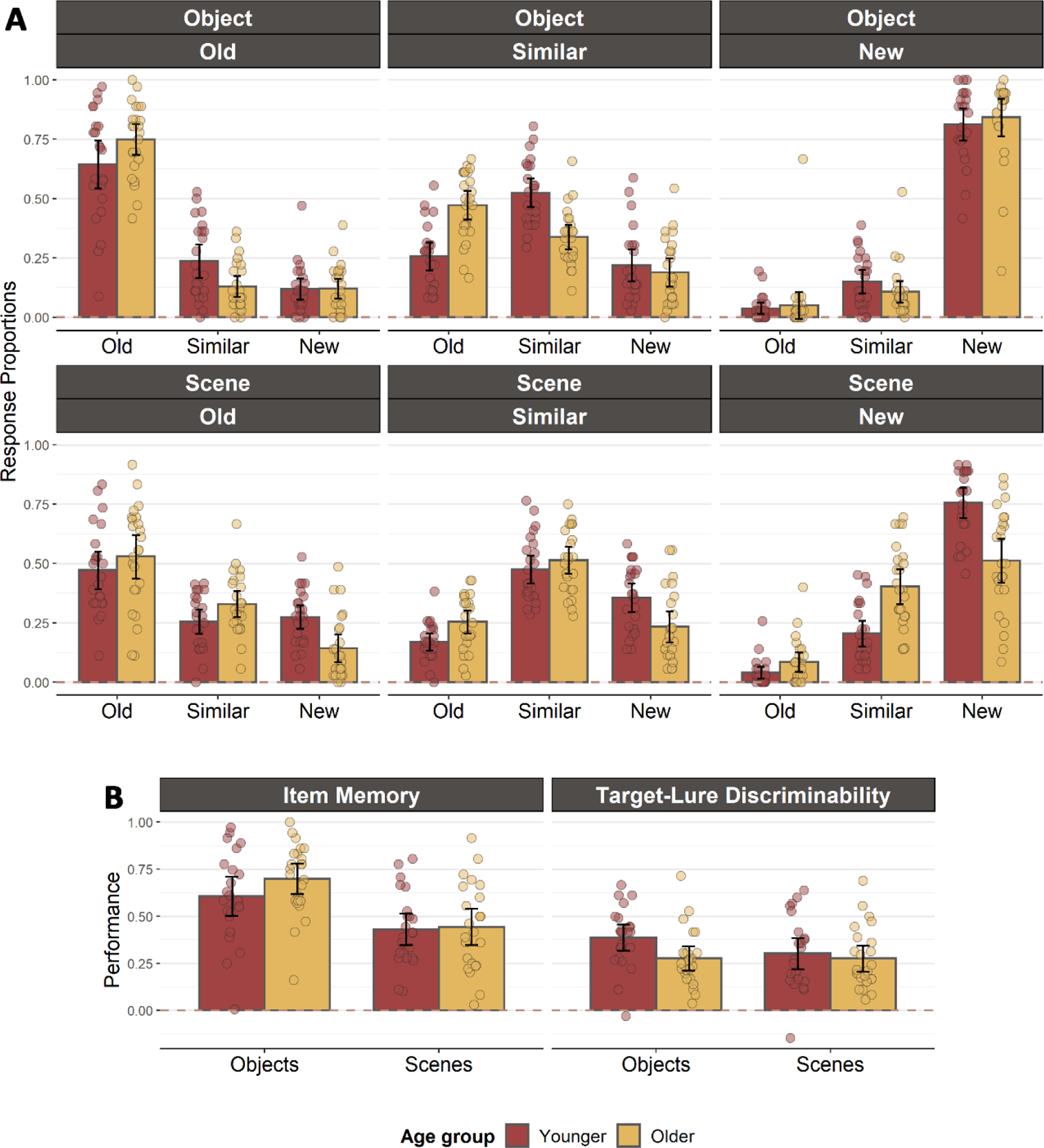
**(A)** Proportion of responses in younger and older adults split separately according to image category, trial type, and response endorsement. **(B)** Item recognition and TLD in younger and older adults.

**Table 2.** Memory performance in younger and older adults. The data reflect the proportion of trials (separately for each image category and trial type) endorsed as old, similar, or new.

| *Image category* | *Younger Adults* |  |  | *Older Adults* |  |  |
| --- | --- | --- | --- | --- | --- | --- |
|  | **Trial type** | **Endorsed as** | **Proportion Mean (SD)** | **Trial type** | **Endorsed as** | **Proportion** |
| ***Scenes*** | Target | Old | 0.47 (0.18) | Target | Old | 0.53 (0.22) |
|  |  | Similar | 0.25 (0.11) |  | Similar | 0.33 (0.13) |
|  |  | New | 0.27 (0.11) |  | New | 0.14 (0.14) |
|  | Lure | Old | 0.17 (0.08) | Lure | Old | 0.25 (0.11) |
|  |  | Similar | 0.47 (0.14) |  | Similar | 0.51 (0.14) |
|  |  | New | 0.36 (0.14) |  | New | 0.23 (0.15) |
|  | New | Old | 0.04 (0.06) | New | Old | 0.08 (0.10) |
|  |  | Similar | 0.20 (0.13) |  | Similar | 0.40 (0.17) |
|  |  | New | 0.76 (0.15) |  | New | 0.51 (0.21) |
| ***Objects*** | Target | Old | 0.64 (0.23) | Target | Old | 0.75 (0.15) |
|  |  | Similar | 0.24 (0.16) |  | Similar | 0.13 (0.10) |
|  |  | New | 0.12 (0.10) |  | New | 0.12 (0.10) |
|  | Lure | Old | 0.26 (0.14) | Lure | Old | 0.47 (0.14) |
|  |  | Similar | 0.52 (0.14) |  | Similar | 0.33 (0.12) |
|  |  | New | 0.21 (0.15) |  | New | 0.19 (0.14) |
|  | New | Old | 0.04 (0.05) | New | Old | 0.05 (0.13) |
|  |  | Similar | 0.15 (0.11) |  | Similar | 0.11 (0.11) |
|  |  | New | 0.81 (0.16) |  | New | 0.84 (0.18) |

The TLD metrics were entered into an analogous 2 (age group) x 2 (image category) mixed effects ANOVA. The ANOVA resulted in a null effect of age group (F_(1, 45)_ = 2.343, p = 0.133, partial-η^2^ = 0.049), while the main effect of category and the age x category interaction were both significant (category: F_(1, 45)_ = 4.741, p = 0.035, partial-η^2^ = 0.095; interaction: F_(1, 45)_ = 4.353, p = 0.043, partial-η^2^ = 0.088). The category effect arose because of better discrimination performance for objects than scenes. The age x category interaction reflected significantly lower discrimination performance in older relative to younger adults for objects (t_(44.44)_ = 2.406, p = 0.020), but a null effect of age for scenes (t_(43.10)_ = 0.509, p = 0.613).

### Category-level PSA results

Similarity indices in the LOC and PPA ROIs revealed reliable category-level effects in younger and older adults for both stimulus categories (see Figure 6A). The category-level similarity indices were entered into a 2 (age group) x 2 (ROI) x 2 (hemisphere) x 2 (image category) mixed effects ANOVA, the outcome of which is reported in Table 3. The ANOVA revealed a main effect of age group, an age x ROI interaction, and an age x category interaction. Significant interaction effects which did not include the factor of age group, and thus are not discussed further, were a significant 2-way interaction between ROI and hemisphere, and a 3-way interaction between ROI, hemisphere, and category. Follow-up analyses of the age x ROI interaction revealed significantly lower category-level similarity in older adults in the PPA (t_(43.60)_ = 3.490, p = 0.001), but no age differences in the LOC (t_(45.99)_ = 0.269, p = 0.789). The age x category interaction was reflective of significant age differences for scenes (t_(42.14)_ = 4.313, p < 0.001), but null effects of age for objects (t_(43.60)_ = 0.416, p = 0.680). Therefore, as is evident in Figure 6A, age-related attenuation of neural differentiation at the category level was observed only in the PPA and only for scene trials.

**Figure 6:**
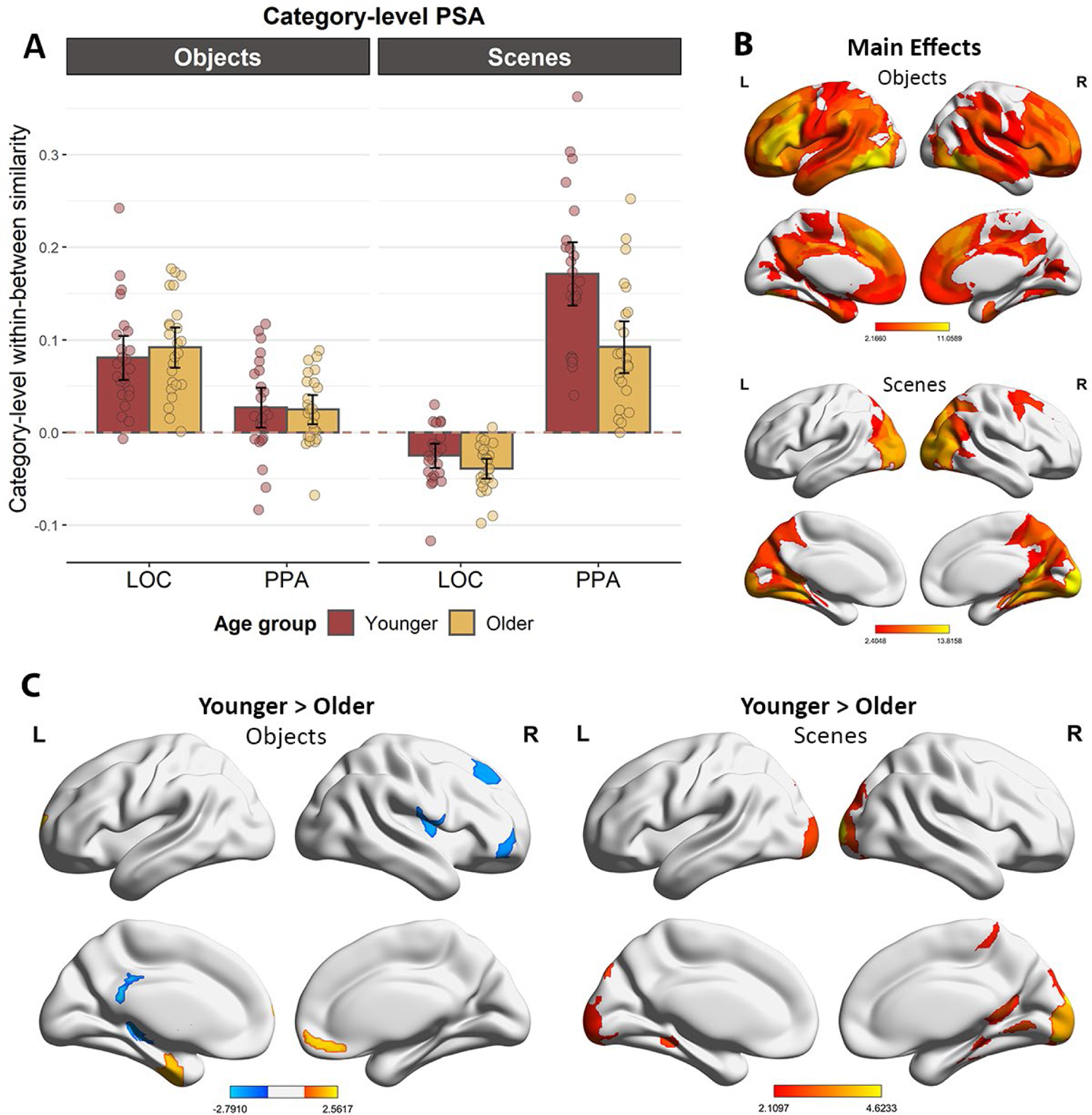
**(A)** Category-level similarity indices in the LOC and PPA ROIs, demonstrating category-level age-related neural dedifferentiation for scenes in the PPA. **(B)** Category-level similarity effects for objects and scenes across age groups. **(C)** Regions exhibiting reliable age differences in neural similarity for scenes and objects (red = greater similarity for younger adults; blue = greater similarity for older adults).

**Table 3:** Outcome of the 2 (age group) x 2 (ROI) x 2 (Hemisphere) x 2 (Category) mixed effects ANOVA performed on the category-level within – between similarity metrics. Significant effects are denoted by *

| | df | F | partial-$\eta^2$ | p-value |
| --- | --- | --- | --- | --- |
| **Age group** | 1,46 | 8.206 | 0.151 | **0.006 \*** |
| **ROI** | 1,46 | 106.438 | 0.698 | **< 0.001 \*** |
| Hemisphere | 1,46 | 0.076 | 0.002 | 0.784 |
| Category | 1,46 | 0.579 | 0.012 | 0.450 |
| **Age x ROI** | 1,46 | 15.412 | 0.251 | **< 0.001 \*** |
| Age x Hemisphere | 1,46 | 0.406 | 0.009 | 0.527 |
| **Age x Category** | 1,46 | 9.947 | 0.178 | **0.003 \*** |
| ROI x Hemisphere | 1,46 | 2.964 | 0.061 | 0.092 |
| **ROI x Category** | 1,46 | 139.133 | 0.752 | **< 0.001 \*** |
| Hemisphere x Category | 1,46 | 0.155 | 0.003 | 0.696 |
| Age x ROI x Hemisphere | 1,46 | < 0.001 | < 0.001 | 0.984 |
| Age x ROI x Category | 1,46 | 1.863 | 0.039 | 0.179 |
| Age x Hemisphere x Category | 1,46 | 0.135 | 0.003 | 0.715 |
| **ROI x Hemisphere x Category** | 1,46 | 9.196 | 0.167 | **0.004 \*** |
| Age x ROI x Hemisphere x Category | 1,46 | 0.304 | 0.007 | 0.584 |

Turning to the whole-brain category-level PSA, Figure 6B depicts those AICHA parcels which exhibited reliable positive scene and object main effects separately in both age groups. Scene-selective effects were primarily evident across the occipital and posterior temporal cortex, the parahippocampal and fusiform gyri, and the retrosplenial complex. By contrast, object-selective effects were more widely distributed, and were prominent in fronto-parietal and anterior temporal cortex. Regions demonstrating reliable age differences in category-level similarity are depicted in Figure 6C. For scene trials, older adults demonstrated lower category-level similarity in the occipital cortex, as well as many of the regions typically implicated in scene processing, such as the parahippocampal cortex and the retrosplenial complex. For object trials, age effects were relatively sparse, with younger adults showing greater similarity than older adults in two clusters in the anterior fusiform gyrus and in the medial prefrontal cortex. However, greater similarity for objects in older adults was evident in a number of frontal regions, along with the posterior hippocampus and posterior cingulate cortex. Therefore, consistent with the ROI analyses, age-related neural dedifferentiation at the category-level was most consistently observed during scene processing in scene-selective cortical regions.

### Item-level PSA results

Analogously to the analyses of category-level PSA, item-level similarity metrics were subjected to a 2 (age group) x 2 (ROI) x 2 (hemisphere) x 2 (image category) x 2 (trial type: repeat/lure) mixed effects ANOVA. The data are illustrated in Figure 7A, and the statistical outcomes of the ANOVA are reported in Table 4. As is evident from the table, the ANOVA revealed a significant main effect of age group, which was driven by greater similarity metrics in younger relative to older adults. Additionally, the ANOVA yielded a main effect of ROI, reflective of greater item similarity in the LOC than in the PPA. The ANOVA also revealed a statistically equivocal category x age interaction (p = 0.050). Follow-up pairwise comparisons indicated that older adults exhibited lower similarity for both objects (t_(45.68)_ = 4.053, p < 0.001) and scenes (t_(45.35)_ = 2.273, p = 0.028), although the age effect was seemingly greater for the objects. We note, however, that the estimation of item-level differentiation for scenes is confounded by differential scene-related ‘baseline’ (between-item) similarity in younger and older adults. Because older adults exhibit relatively lower similarity across all scene stimuli (see category-level results), age differences in scene selectivity might have been harder to detect at the item-level. Effects which did not interact with the factor of age group, and thus are not discussed further, included a significant 2-way interaction between ROI and hemisphere and a 4-way interaction between ROI, hemisphere, category, and trial type. In summary, these results revealed that, in contrast to the category-level PSA, age-related neural dedifferentiation was evident at the item level for both scene and object stimuli, ostensibly to a greater extent for objects. Moreover, the analyses revealed that neural similarity was lower in older adults regardless of trial type (repeat versus lure). Indeed, with the exception of the four-way interaction, effects of trial type were uniformly absent.

**Figure 7:**
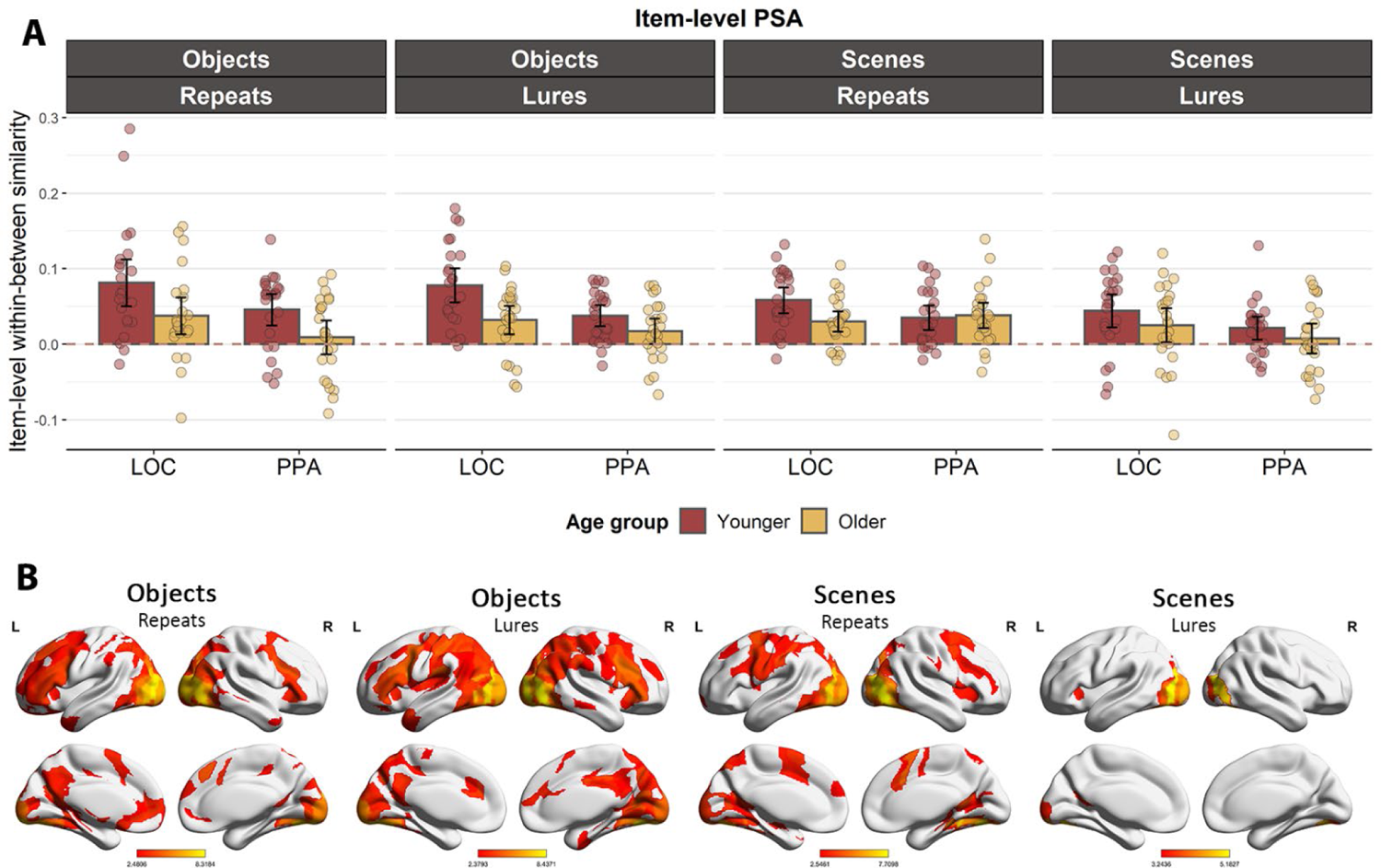
**(A)** Item-level similarity indices in the LOC and PPA ROIs, revealing reduced similarity in older adults for objects and scenes across both repeats and lures. **(B)** Reliable item-level effects for objects and scenes in both age groups.

**Table 4:** Outcome of the 2 (age group) x 2 (ROI) x 2 (Hemisphere) x 2 (Category) x 2 (Trial type) mixed effects ANOVA performed on the item-level within – between similarity metrics. Significant effects are denoted by *

|  | *df* | *F* | *partial-η<sup>2</sup>* | *p-value* |
| --- | --- | --- | --- | --- |
| **Age group** | **1, 46** | **20.915** | **0.313** | **< 0.001 \*** |
| **ROI** | **1, 46** | **21.647** | **0.32** | **< 0.001 \*** |
| Hemisphere | 1, 46 | 1.836 | 0.038 | 0.182 |
| Category | 1, 46 | 3.249 | 0.066 | 0.078 |
| Trial Type | 1, 46 | 2.136 | 0.044 | 0.151 |
| Age x ROI | 1, 46 | 3.404 | 0.069 | 0.071 |
| Age x Hemisphere | 1, 46 | 0.005 | < 0.001 | 0.945 |
| **Age x Category** | **1, 46** | **4.043** | **0.081** | **0.050 \*** |
| Age x Trial Type | 1, 46 | 0.015 | < 0.001 | 0.903 |
| ROI x Hemisphere | 1, 46 | 2.359 | 0.049 | 0.131 |
| **ROI x Category** | **1, 46** | **4.89** | **0.096** | **0.032 \*** |
| Hemisphere x Category | 1, 46 | 0.188 | 0.004 | 0.667 |
| ROI x Type | 1, 46 | 0.211 | 0.005 | 0.648 |
| Hemisphere x Type | 1, 46 | 1.142 | 0.024 | 0.291 |
| Category x Type | 1, 46 | 1.976 | 0.041 | 0.166 |
| Age x ROI x Hemisphere | 1, 46 | 0.012 | < 0.001 | 0.913 |
| Age x ROI x Category | 1, 46 | 0.018 | < 0.001 | 0.894 |
| Age x Hemisphere x Category | 1, 46 | 0.813 | 0.017 | 0.372 |
| Age x ROI x Type | 1, 46 | 0.043 | 0.001 | 0.837 |
| Age x Hemisphere x Type | 1, 46 | 0.002 | < 0.001 | 0.960 |
| Age x Category x Type | 1, 46 | 0.304 | 0.007 | 0.584 |
| ROI x Hemisphere x Category | 1, 46 | 0.015 | < 0.001 | 0.905 |
| ROI x Hemisphere x Type | 1, 46 | 0.027 | 0.001 | 0.870 |
| ROI x Category x Type | 1, 46 | 1.41 | 0.03 | 0.241 |
| Hemisphere x Category x Type | 1, 46 | 1.735 | 0.036 | 0.194 |
| Age x ROI x Hemisphere x Category | 1, 46 | 0.713 | 0.015 | 0.403 |
| Age x ROI x Hemisphere x Type | 1, 46 | 2.137 | 0.044 | 0.151 |
| Age x ROI x Category x Type | 1, 46 | 2.368 | 0.049 | 0.131 |
| Age x Hemisphere x Category x Type | 1, 46 | 0.268 | 0.006 | 0.607 |
| **ROI x Hemisphere x Category x Type** | **1, 46** | **9.052** | **0.164** | **0.004 \*** |
| Age x ROI x Hemisphere x Category x Type | 1, 46 | 0.901 | 0.019 | 0.348 |

The outcomes of the exploratory whole brain item-level PSAs are illustrated in Figure 7B. In all cases, reliable across-participant item-level effects were observed across much of the occipital cortex. However, whereas effects for scene lures were largely restricted to these occipital regions, effects for scene repeats and for objects (repeats and lures alike) extended into parietal and the posterior temporal cortex (e.g., parahippocampal and fusiform gyri). Additionally, each of these trial types exhibited robust effects in parietal and frontal cortex. The outcomes of contrasts between age groups were hard to interpret. Age differences were evident in only a few AICHA parcels, which were scattered across the cortex with little clustering across neighboring parcels. One notable exception was evidence for greater item-level similarity in younger adults for object repeats and lures in posterior parahippocampal cortex and the lateral occipital areas, a result which corresponded with our ROI-based analyses. The figures summarizing the age-group contrasts are available from the 1^st^ author upon request.

### Relationship between neural differentiation and memory performance

Motivated by prior reports of a positive, age-invariant relationship between neural differentiation in the PPA and memory performance (e.g., Koen et al., 2019; Srokova et al., 2020), we performed multiple regression analyses to examine the relationship between TLD and neural similarity. The TLD scores were highly correlated between objects and scenes (r_partial_ = 0.670, p < 0.001, controlling for age group). Therefore, to reduce the number of multiple comparisons we averaged the scores across categories to produce a single mean TLD metric. The analysis approach consisted of two steps. First, a regression model was defined using TLD as the dependent variable, and the variables of age group, within-between similarity, and their interaction term as the predictors. A statistically significant age group x similarity interaction would suggest that the relationship between TLD and similarity is moderated by age group, motivating follow-up analyses in the form of the computation of zero-order correlations separately for each age group. A non-significant interaction term would indicate that any relationship between the neural and behavioral variables was age-invariant. In these cases, the relationship was quantified by the partial correlation between neural similarity and TLD, controlling for age group. We performed a total of 4 regression analyses, separately for object-related similarity in the LOC and scene-related similarity in the PPA, operationalized either at the item or category level.

Turning first to category-level differentiation of scene stimuli, the age group x scene similarity interaction in the PPA was not significant (p = 0.166), indicating that any potential relationship between scene-related similarity and TLD is not moderated by age group. A partial correlation (controlling for age group) between scene-related similarity in the PPA and the TLD was statistically significant (r_partial_ = 0.390, p = 0.007; Figure 8), revealing evidence for an age-invariant relationship. Turning to category-level differentiation for objects, the interaction between age group and object similarity in the LOC was also non-significant (p = 0.393), as however was the partial correlation between LOC similarity and TLD (r*_partial_* = 0.277, p = 0.062). In conclusion, our analyses demonstrate that the relationship between TLD and category-level similarity was restricted to scene-related similarity in the PPA. The regression and correlation analyses examining the association between TLD and neural similarity at the item-level (either for repeats or lures) did not yield any statistically significant relationships.

**Figure 8:**
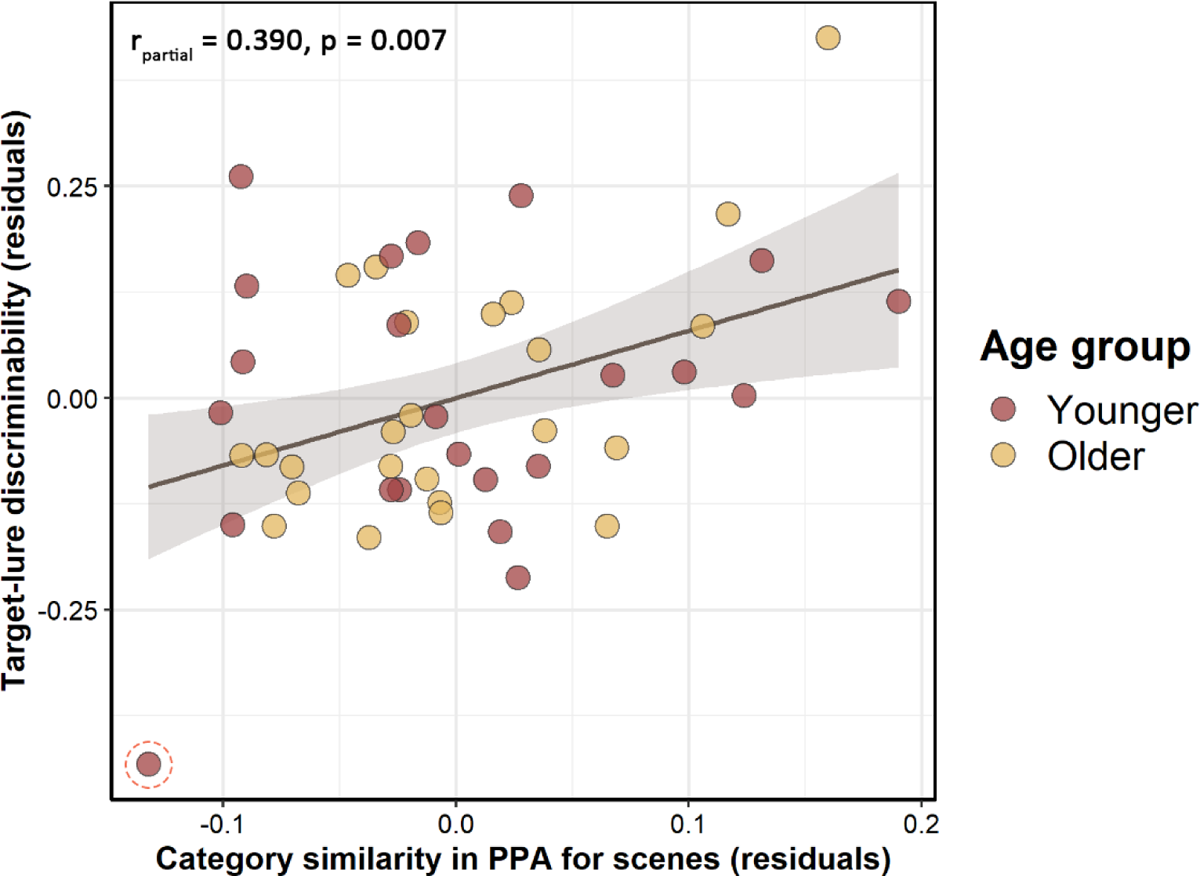
Age-invariant relationship between mean TLD and scene-related category-level neural differentiation in the PPA. The relationship remains significant (p = 0.035) following exclusion of the highlighted outlier.

Lastly, we performed partial correlations (controlling for age) to examine the relationships between the category-level and item-level metrics of neural differentiation (for the item-level metrics, we collapsed similarity across exact repeats and lures). Category- and item-level metrics of both scene and object differentiation were uncorrelated in both ROIs (p min. = 0.148).

## Discussion

In the present study, we examined the effects of age on object- and scene-related neural differentiation at the levels of image categories and individual stimulus exemplars. Consistent with prior reports (e.g., Koen et al., 2019), multi-voxel pattern similarity analyses at the category-level identified lower differentiation in the older relative to the younger age group in scene-selective, but not object-selective, cortical regions. In contrast, item-level analyses revealed reduced neural similarity in older adults for exemplars of both stimulus categories and regardless of trial type (repeat vs. similar lure). Also consistent with prior findings, regression analyses revealed that greater category-level scene selectivity in the PPA was predictive of higher TLD regardless of age group. Crucially, the present study provides novel evidence that null age effects in category-level differentiation do not necessarily extend to metrics of item-level neural differentiation, suggesting that these two measures of neural selectivity reflect distinct neural mechanisms.

### Behavioral Results

Participants completed the retrieval phase of the MST outside of the scanner immediately following the scanned encoding phase. We found no age differences in item recognition, consistent with numerous prior findings of relatively preserved familiarity-based recognition memory in older age (for review, see Koen and Yonelinas, 2014). In contrast, object, but not scene TLD was lower in older adults, indicative of an age-related decline in the ability to discriminate between previously viewed images of objects and their similar lures. These findings are consistent with prior reports, which indicate that age differences in lure discriminability are frequently observed in MST variants utilizing object stimuli (see Stark et al., 2019), while age differences are less common or weaker in variants of the task employing spatial or scene stimuli (Reagh et al., 2016; Stark & Stark, 2017; Berron et al., 2018; note that, in most prior studies, lure discriminability was measured using an alternate metric). This object-specific decline in putative behavioral measures of pattern separation has previously been attributed to age-related decline in the functional integrity of perirhinal and lateral entorhinal cortex, considering that the regions have been implicated in object processing and, additionally, are a major source of object information flowing into the hippocampus (Reagh et al., 2016). However, the notion that scene-specific input to the hippocampus is relatively preserved with increasing age is difficult to reconcile with present and prior findings of robust neural dedifferentiation in scene-selective cortical regions, including the posterior parahippocampal cortex (which forms part of the PPA). We conjecture that the age differences in TLD for objects, but not scenes, may be reflective of a greater conceptual confusability between similar objects. Relative to scenes, object stimuli contain less perceptual information diagnostic for the discrimination between conceptually identical items. Therefore, successful discrimination between similar objects may require relatively greater mnemonic precision at the perceptual level. Given the recent evidence suggestive of an age-related decrease in mnemonic precision (Nilakantan et al., 2018; Korkki et al., 2020), along with an increased reliance on gist-based and conceptual information (Srokova et al., 2022), older adults may have greater difficulty discriminating between objects due a relative lack of diagnostic perceptual information.

### Effects of age on category-level neural differentiation

Whereas age-related neural dedifferentiation has consistently been reported for scene images in scene-selective cortical regions, neural dedifferentiation for objects, words, or faces in their respective category-selective regions has been identified much less consistently (for review, see Koen & Rugg, 2019; Koen et al., 2020). The mechanisms responsible for category-level neural dedifferentiation and the reasons for its seeming regional-specificity are the subjects of debate. One candidate mechanism, proposed by Li and colleagues (2001), posits that age-related neural dedifferentiation reflects a decline with age in the integrity of the ascending dopaminergic neuromodulatory system. From this perspective, reductions in selectivity are caused by reduced dopaminergic availability, which compromises neural signal-to-noise ratio, and hence the fidelity of neural representation. Somewhat analogously, neural dedifferentiation has also been hypothesized to reflect age-related decline in γ-aminobutyric acid (GABA) inhibitory neurotransmission (Lalwani et al. 2019; Cassady et al., 2019, 2020; Chamberlain et al., 2021) and a consequent broadening of neural tuning. However, the current and prior evidence for regional-specificity of neural dedifferentiation (at least at the category level) arguably challenges the notion that the phenomenon is attributable such factors, at least if they are considered to operate cortex-wide.

A possible explanation for the ubiquity of category-level age-related dedifferentiation of scene stimuli stems from findings that the magnitude of fMRI BOLD responses in scene-selective cortical areas, such as the PPA, are strongly modulated by perceptual complexity and attentional factors (Aminoff et al., 2013). As we have proposed previously (Srokova et al., 2020), age-related decline in neural selectivity for scenes might be a consequence of decline in the ability to process complex visual inputs and, especially, difficulty in differentiating the multiple, spatially distributed elements that constitute a scene. Alternately, or additionally, scene dedifferentiation might be related to reduced availability of domain-general attentional resources (see Serences et al., 2004; Gazzaley et al., 2005), such that older adults fail to exert sufficient attentional control over scene processing (Bouhassoun et al., 2022). These potential mechanisms assume that the null effects of age for faces and object selectivity that have been reported reflect their relatively lower visual complexity and lower attentional demands. As we discuss below, on its face, this account is compromised by the present finding that age effects on item-level selectivity extend beyond scenes to include objects.

### Effects of age on item-level neural differentiation

As noted in the Introduction, it might be assumed that age-related category-level neural dedifferentiation is driven by a decline in the selectivity of neural patterns elicited by individual category exemplars. However, no prior study has directly examined whether age differences in category-level differentiation are associated with age differences at the item-level. As outlined in the introduction, Goh et al. (2010) reported that while younger adults exhibited robust univariate repetition suppression effects for exact repeats of faces, older adults exhibited effects for both repeats and perceptually similar lures, a finding indicative of item-level dedifferentiation. Additionally, in two studies that employed MVPA, it was reported that pattern similarity for exact repetitions was lower in older adults (Bowman et al., 2019; Trelle et al., 2019). By contrast, two other studies reported null effects of age on the neural similarity between successive stimulus presentations (St-Laurent et al., 2019; Zheng et al., 2018). The present findings are consistent with those prior studies that reported a decline with age in item-level neural differentiation.

The present study offers novel evidence opposing the idea that category- and item-level differentiation depend on the same neural mechanisms. We demonstrate that absent age effects on object selectivity at the category level do not extend to neural selectivity at the level of individual objects, at least as this is indexed by multi-voxel PSA. Rather, age-related dedifferentiation at the item level was evident for both scenes and objects and, if anything, was greater for objects. In attempting to understand the implications of this dissociation between category- and item-level metrics, we note that metrics of item-level differentiation quantify the extent to which a given item elicits neural patterns that are *distinct* from those elicited by other items belonging to the same category (i.e., item-specific information). In contrast, category-level similarity indexes the extent to which neural patterns are *shared* between category exemplars, potentially reflecting processes that are engaged by all (or most) members of a category. There is no obvious reason why these two expressions of neural selectivity should be related to one another. Indeed, consistent with this point, and regardless of age group or stimulus category, item- and category-level similarity metrics were uncorrelated across participants in the present study.

Lastly, we note that in both age groups, we were unable to identify any effect on item-level selectivity of the ‘repeat’ vs. ‘similar lure’ manipulation. We have no ready explanation for this null finding, not least since our participants demonstrated the ability to discriminate between repeats and lures on the post-scan memory test; our expectation was that, at the least, we would find evidence for attenuated selectivity for the lures in the younger age group (cf. Goh et al., 2010). One possibility is that the failure to detect differences in neural selectivity between the two classes of items reflects a mismatch between the spatial resolution at which these differences were expressed in the cortex and the resolution of our imaging methods.

### Relationship between category-level neural differentiation and memory performance

Category-level neural selectivity has consistently been reported to correlate positively across participants with memory performance in an age-invariant manner (Koen et al., 2019; Srokova et al., 2020). On the basis of the limited available data (the majority of prior studies where a relationship between neural selectivity and memory performance was reported collapsed across different category-selective regions and exemplars), this relationship appears to be selective for metrics of scene selectivity derived from the PPA. The present findings are fully consistent with these prior results. Why PPA selectivity, but seemingly not selectivity metrics derived from other regions, should be sensitive to memory performance is currently unclear (for further discussion, see Srokova et al., 2022).

## Conclusions

In summary, our data replicate and extend prior studies of age-related dedifferentiation. First, we find that, at the category level, evidence for neural dedifferentiation is limited to scene-selective cortical regions. Crucially, however, this dissociation does not extend to neural differentiation at the item level, when age-related dedifferentiation was evident for both scene and object images. Thus, item- and category-level metrics of neural differentiation are differentially sensitive to increasing age, and likely reflect distinct neural mechanisms. Of importance, the consistently reported associations between neural differentiation, age and cognitive performance should motivate future research to investigate its mechanistic underpinnings.

## Acknowledgements

This work was supported by the National Institute of Aging Grants R56AG068149 and RF1AG039103 and an award from BvB Dallas. The authors would like to acknowledge Joshua Olivier, Nehal Shahanawaz, and Eduardo Hernandez for their assistance with recruitment and neuropsychological assessments.

## Notes

### Competing Interest Statement

The authors have declared no competing interest.

## References

Aminoff, E. M., Kveraga, K., & Bar, M. (2013). The role of the parahippocampal cortex in cognition. Trends in cognitive sciences, 17(8), 379–390.

Bailey, I. L., & Lovie-Kitchin, J. E. (2013). Visual acuity testing. From the laboratory to the clinic. Vision research, 90, 2–9.

Barron, H. C., Garvert, M. M., & Behrens, T. E. (2016). Repetition suppression: a means to index neural representations using BOLD?. Philosophical Transactions of the Royal Society B: Biological Sciences, 371(1705), 20150355.

Berron, D., Neumann, K., Maass, A., Schütze, H., Fliessbach, K., Kiven, V., … & Düzel, E. (2018). Age-related functional changes in domain-specific medial temporal lobe pathways. Neurobiology of aging, 65, 86–97.

Bouhassoun, S., Poirel, N., Hamlin, N., & Doucet, G. E. (2022). The forest, the trees, and the leaves across adulthood: Age-related changes on a visual search task containing three-level hierarchical stimuli. Attention, Perception, & Psychophysics, 1-12.

Bowman, C. R., Chamberlain, J. D., & Dennis, N. A. (2019). Sensory representations supporting memory specificity: Age effects on behavioral and neural discriminability. Journal of Neuroscience, 39(12), 2265–2275.

Carp, J., Park, J., Polk, T. A., & Park, D. C. (2011). Age differences in neural distinctiveness revealed by multi-voxel pattern analysis. Neuroimage, 56(2), 736–743.

Cassady, K., Gagnon, H., Freiburger, E., Lalwani, P., Simmonite, M., Park, D. C., … & Polk, T. A. (2020). Network segregation varies with neural distinctiveness in sensorimotor cortex. NeuroImage, 212, 116663.

Cassady, K., Gagnon, H., Lalwani, P., Simmonite, M., Foerster, B., Park, D., … & Polk, T. A. (2019). Sensorimotor network segregation declines with age and is linked to GABA and to sensorimotor performance. Neuroimage, 186, 234–244.

Chamberlain, J. D., Gagnon, H., Lalwani, P., Cassady, K. E., Simmonite, M., Seidler, R. D., … & Polk, T. A. (2021). GABA levels in ventral visual cortex decline with age and are associated with neural distinctiveness. Neurobiology of aging, 102, 170–177.

Chee, M. W., Goh, J. O., Venkatraman, V., Tan, J. C., Gutchess, A., Sutton, B., … & Park, D. (2006). Age-related changes in object processing and contextual binding revealed using fMR adaptation. Journal of cognitive neuroscience, 18(4), 495–507.

Ferris III, Frederick L., et al. “New visual acuity charts for clinical research.” American journal of ophthalmology 94.1 (1982): 91–96.

Gazzaley, A., Cooney, J. W., McEvoy, K., Knight, R. T., & D’esposito, M. (2005). Top-down enhancement and suppression of the magnitude and speed of neural activity. Journal of cognitive neuroscience, 17(3), 507–517.

Goh, J. O., Suzuki, A., & Park, D. C. (2010). Reduced neural selectivity increases fMRI adaptation with age during face discrimination. Neuroimage, 51(1), 336–344.

Joliot, M., Jobard, G., Naveau, M., Delcroix, N., Petit, L., Zago, L., … & Tzourio-Mazoyer, N. (2015). AICHA: An atlas of intrinsic connectivity of homotopic areas. Journal of neuroscience methods, 254, 46–59.

Koen, J. D., & Rugg, M. D. (2019). Neural dedifferentiation in the aging brain. Trends in Cognitive Sciences, 23(7), 547–559.

Koen, J. D., & Yonelinas, A. P. (2014). The effects of healthy aging, amnestic mild cognitive impairment, and Alzheimer’s disease on recollection and familiarity: A meta-analytic review. Neuropsychology review, 24, 332–354.

Koen, J. D., Hauck, N., & Rugg, M. D. (2019). The relationship between age, neural differentiation, and memory performance. Journal of Neuroscience, 39(1), 149–162.

Koen, J. D., Srokova, S., & Rugg, M. D. (2020). Age-related neural dedifferentiation and cognition. Current Opinion in Behavioral Sciences, 32, 7–14.

Korkki, S. M., Richter, F. R., Jeyarathnarajah, P., & Simons, J. S. (2020). Healthy ageing reduces the precision of episodic memory retrieval. Psychology and Aging, 35(1), 124.

Kriegeskorte, N., Mur, M., & Bandettini, P. A. (2008). Representational similarity analysis-connecting the branches of systems neuroscience. Frontiers in systems neuroscience, 4.

Lalwani, P., Gagnon, H., Cassady, K., Simmonite, M., Peltier, S., Seidler, R. D., … & Polk, T. A. (2019). Neural distinctiveness declines with age in auditory cortex and is associated with auditory GABA levels. Neuroimage, 201, 116033.

Li, S. C., Lindenberger, U., & Sikström, S. (2001). Aging cognition: from neuromodulation to representation. Trends in cognitive sciences, 5(11), 479–486.

Maass, A., Berron, D., Harrison, T. M., Adams, J. N., La Joie, R., Baker, S., … & Jagust, W. J. (2019). Alzheimer’s pathology targets distinct memory networks in the ageing brain. brain, 142(8), 2492–2509.

Mumford, J. A., Davis, T., & Poldrack, R. A. (2014). The impact of study design on pattern estimation for single-trial multivariate pattern analysis. Neuroimage, 103, 130–138.

Nilakantan, A. S., Bridge, D. J., VanHaerents, S., & Voss, J. L. (2018). Distinguishing the precision of spatial recollection from its success: Evidence from healthy aging and unilateral mesial temporal lobe resection. Neuropsychologia, 119, 101–106.

Park, D. C., Polk, T. A., Park, R., Minear, M., Savage, A., & Smith, M. R. (2004). Aging reduces neural specialization in ventral visual cortex. Proceedings of the National Academy of Sciences, 101(35), 13091–13095.

Park, J., Carp, J., Hebrank, A., Park, D. C., & Polk, T. A. (2010). Neural specificity predicts fluid processing ability in older adults. Journal of Neuroscience, 30(27), 9253–9259.

Park, J., Carp, J., Kennedy, K. M., Rodrigue, K. M., Bischof, G. N., Huang, C. M., … & Park, D. C. (2012). Neural broadening or neural attenuation? Investigating age-related dedifferentiation in the face network in a large lifespan sample. Journal of Neuroscience, 32(6), 2154–2158.

Payer, D., Marshuetz, C., Sutton, B., Hebrank, A., Welsh, R. C., & Park, D. C. (2006). Decreased neural specialization in old adults on a working memory task. Neuroreport, 17(5), 487–491.

Peirce, J., Gray, J. R., Simpson, S., MacAskill, M., Höchenberger, R., Sogo, H., … & Lindeløv, J. K. (2019). PsychoPy2: Experiments in behavior made easy. Behavior research methods, 51, 195–203.

R Core Team (2020) R: a language and environment for statistical computing. Vienna: R Foundation.

Raven, J., Raven, J.C., Courth, J.H. (2000) The advanced progressive matrices. In: Manual for Raven’s progressive matrices and vocabulary scales, Section 4. San Antonio: Harcourt Assessment.

Reagh, Z. M., Ho, H. D., Leal, S. L., Noche, J. A., Chun, A., Murray, E. A., & Yassa, M. A. (2016). Greater loss of object than spatial mnemonic discrimination in aged adults. Hippocampus, 26(4), 417–422.

Reitan, R.M., Wolfson, D. (1985) The Halstead-Reitan neuropsychological test battery: therapy and clinical interpretation. Tucson: Neuropsychological.

Rissman, J., Gazzaley, A., & D’Esposito, M. (2004). Measuring functional connectivity during distinct stages of a cognitive task. Neuroimage, 23(2), 752–763.

Serences, J. T., Schwarzbach, J., Courtney, S. M., Golay, X., & Yantis, S. (2004). Control of object-based attention in human cortex. Cerebral cortex, 14(12), 1346–1357.

Smith, A. (1982) Symbol digit modalities test (SDMT) manual. Los Angeles: Western Psychological Services.

Sommer, V. R., Fandakova, Y., Grandy, T. H., Shing, Y. L., Werkle-Bergner, M., & Sander, M. C. (2019). Neural pattern similarity differentially relates to memory performance in younger and older adults. Journal of Neuroscience, 39(41), 8089–8099.

Spreen O., Benton A.L. (1977) Neurosensory center comprehensive examination for aphasia. Victoria: Neuropsychology Laboratory.

Srokova, S., Hill, P. F., & Rugg, M. D. (2022). The retrieval-related anterior shift is moderated by age and correlates with memory performance. Journal of Neuroscience, 42(9), 1765–1776.

Srokova, S., Hill, P. F., Koen, J. D., King, D. R., & Rugg, M. D. (2020). Neural differentiation is moderated by age in scene-selective, but not face-selective, cortical regions. ENeuro, 7(3).

Stark, S. M., & Stark, C. E. (2017). Age-related deficits in the mnemonic similarity task for objects and scenes. Behavioural brain research, 333, 109–117.

Stark, S. M., Kirwan, C. B., & Stark, C. E. (2019). Mnemonic similarity task: A tool for assessing hippocampal integrity. Trends in cognitive sciences, 23(11), 938–951.

St-Laurent, M., & Buchsbaum, B. R. (2019). How multiple retrievals affect neural reactivation in young and older adults. The Journals of Gerontology: Series B, 74(7), 1086–1100.

Trelle, A. N., Henson, R. N., & Simons, J. S. (2019). Neural evidence for age-related differences in representational quality and strategic retrieval processes. Neurobiology of Aging, 84, 50–60.

Voss, M. W., Erickson, K. I., Chaddock, L., Prakash, R. S., Colcombe, S. J., Morris, K. S., … & Kramer, A. F. (2008). Dedifferentiation in the visual cortex: an fMRI investigation of individual differences in older adults. Brain research, 1244, 121–131.

Wechsler, D. (1981) WAIS-R: Wechsler adult intelligence scale-revised. New York: The Psychological Corporation.

Wechsler, D. (2009) Wechsler memory scale, Ed 4. San Antonio: The Psychological Corporation.

Wechsler, D. (2011) The Test of Premorbid Function (TOPF). The Psychological Corporation, San Antonio, TX

Yassa, M. A., Mattfeld, A. T., Stark, S. M., & Stark, C. E. (2011). Age-related memory deficits linked to circuit-specific disruptions in the hippocampus. Proceedings of the National Academy of Sciences, 108(21), 8873–8878.

Zebrowitz, L., Ward, N., Boshyan, J., Gutchess, A., & Hadjikhani, N. (2016). Dedifferentiated face processing in older adults is linked to lower resting state metabolic activity in fusiform face area. Brain research, 1644, 22–31.

Zheng, L., Gao, Z., Xiao, X., Ye, Z., Chen, C., & Xue, G. (2018). Reduced fidelity of neural representation underlies episodic memory decline in normal aging. Cerebral Cortex, 28(7), 2283–2296.

